# Resolving the heterogeneity of diaphragmatic mesenchyme: a novel mouse model of congenital diaphragmatic hernia

**DOI:** 10.1101/2020.05.19.104703

**Authors:** Louise Cleal, Ofelia Martinez-Estrada, You-Ying Chau

## Abstract

Congenital diaphragmatic hernia (CDH) is a relatively common developmental defect with considerable mortality and morbidity. Diaphragm formation is a complex process, involving several cell types, each with different developmental origins. Due to this complexity, the aetiology of CDH is not well understood. The pleuroperitoneal folds (PPFs) and the post hepatic mesenchymal plate (PHMP) are transient structures that are essential during diaphragm development. Using several mouse models including lineage tracing, we demonstrate the heterogeneous nature of the cells that make up the PPFs. The conditional deletion of *Wt1* (Wilms’ Tumour gene) in the non-muscle mesenchyme of the PPFs results in CDH. We show that the fusion of the PPFs and the PHMP to form a continuous band of tissue involves migration of cells from both sources. The PPFs of mutant mice fail to fuse with the PHMP and exhibit increased RALDH2 expression. However, no changes in the expression of genes implicated in epithelial-to-mesenchymal transition (EMT) are observed. Additionally, the mutant PPFs lack migrating myoblasts and muscle connective tissue fibroblasts (TCF4+/GATA4+), suggesting possible interactions between these cell types. Our study demonstrates the importance of the non-muscle mesenchyme in diaphragm development.

**Author Summary:** Congenital diaphragmatic hernia (CDH) is a frequent developmental defect and it remains one of the most difficult problems of perinatology. The defect can be repaired by surgery but it is often associated with complications and total mortalities are still high (50-60%). The causes of CDH are largely unknown. Body cavity formation is a carefully regulated process, and diaphragm formation defines the thoracic and abdominal cavities. Connective tissue and muscle fibres are the known major players involved in diaphragm formation. Our current study emphasises another player, the mesenchymal cells. We manipulate the expression of an important developmental regulator, *Wt1* (Wilms’ tumour gene), in mesenchymal cells in a tissue specific manner using transgenic mouse models. We found that mutant mice can survive till birth, develop diaphragmatic hernia, and die shortly after birth. Ablating *Wt1* in the mesenchymal cells leads to ceased movement, and the failure to form a continuous band of tissue. In our model, disruption of mesenchymal cell movement leads to the attenuation of migration of connective tissue fibroblasts and myoblasts, suggesting possible interactions between these cell types.

## Introduction

Congenital diaphragmatic hernia (CDH) is a severe developmental defect that affects approximately 1:3000 live births^1^ and can have devastating clinical outcomes. The aetiology of CDH is not well understood due to the complexity of diaphragm formation. The mature diaphragm is composed of several tissues including muscle connective tissue, muscle, tendons, nerves, blood vessels, lymphatics and mesothelium^2^. Several embryonic structures are implicated in diaphragm development: the pleuroperitoneal folds (PPFs), the septum transversum (ST), the post hepatic mesenchymal plate (PHMP), and the somites^3,4^. The ST is a thin layer of mesodermal cells overlaying the liver and is formed at approximately E8.5 in the mouse. This is followed by the formation of the PPFs which develop at E10.5-E12.5. A recent study demonstrated that the fibroblasts of PPFs play a crucial role in guiding the expansion and movement of the myogenic cells that originate in the somites and are essential for the correct formation of the muscular component of the diaphragm^5^. The PHMP first appears at around E10.5, and is thought to be derived from the mesenchymal population of the lateral ST^6,4^. The cells of PPFs fuse with the PHMP to form a membranous continuum, separating the thoracic and peritoneal cavities. Subsequently, myoblasts migrate from the somites leading to the muscularisation of this membrane to form the mature diaphragm.

Mutations in *WT1* have been described in patients with CDH^7,8,9,10^. Homozygous *Wt1* null mouse embryos also develop diaphragmatic hernias^11^. During diaphragm development in the mouse, WT1 is expressed in the PPFs, PHMP, ST, mesothelium, and lateral wall body mesenchyme^4^. Mesenchymal cells are present throughout the diaphragm but their origins and cell types are not well defined or understood. One mesenchymal cell population: connective tissue fibroblasts (for which GATA4 and TCF4 are the best markers), is crucial for guiding the migration of myoblasts during diaphragm development, as shown by the conditional deletion of *Gata4* using the Prx1-Cre mouse model^5^. However, the TCF4/GATA4 expressing connective tissue fibroblast population does not substantially overlap with the WT1 expressing nonmuscle mesenchyme in the diaphragm^12^.

To delineate the heterogeneity of the ill-defined mesenchymal cells in the diaphragm, we generated a mouse model in which *Wt1* was conditionally deleted in the Prx1-Cre lineage. In this model, mutant embryos can survive *in utero* but die shortly after birth, which we believe is due to the formation of diaphragmatic hernias. In addition to the CDH phenotype, we show that the developmental origin(s) of the non-muscle mesenchymal cells in the PPF is different from those in the PHMP. Moreover, we show data providing cellular insights into the migration of the PPFs during the formation of the diaphragm.

## Results

### Diaphragm development is disrupted in *Prx1^Cre/+^;Wt1^loxp/loxp^* embryos

In our model, male *Prx1^Cre/+^;Wt1^loxp/+^* mice were crossed with female *Wt1^loxp/loxp^* mice to conditionally inactivate *Wt1* using Prx1-Cre. No live mutants (*Prx1^Cre/+^;Wt1^loxp/loxp^*) were present when the litters were genotyped at approximately 2-3 weeks old (n>50). Given the important role of Wt1 in regulating key developmental processes^13^, we suspected the phenotypes of the mutants likely resulted in embryonic lethality. However, mutant embryos appeared to be grossly normal (externally) at all stages analysed (E11.5, E12.5, E14.5, E16.5, E18.5 and E19.5). The number of mutant embryos obtained at each stage is summarised in Table1. When the pregnant dams were left to give birth, it was apparent that mutant pups were born alive but died within a few hours. Obtaining mutant mice that survived until birth led us to hypothesise that their death may have been caused by an inability to breath. Diaphragmatic defects typically result in disrupted breathing^1^. As mentioned previously, *Wt1* null mouse embryos also develop diaphragmatic hernias^11^. Therefore, we hypothesised that the *Prx1^Cre/+^;Wt1^loxp/loxp^* embryos may have diaphragmatic hernias.

**Table 1.** The number of the (Prx1^Cre/+^;Wt1^loxP/loxp^) embryos obtained at each developmental stage, compared with the expected number.

| EMBRYO STAGE | TOTAL # EMBRYOS | # EXPECTED MUTANTS | # ACTUAL MUTANTS |
| --- | --- | --- | --- |
| E10.5 | 27 | 6.75 (25%) | 6 (22%) |
| E11.5 | 18 | 4.5 (25%) | 4 (22%) |
| E12.5 | 26 | 6.5 (25%) | 6 (23%) |
| E14.5 | 9 | 2.25 (25%) | 3 (33%) |
| E16.5 | 22 | 5.5 (25%) | 8 (36%) |
| E18.5 | 6 | 1.5 (25%) | 1 (17%) |
| E19.5 | 25 | 6.25 (25%) | 4 (16%) |
| P0 | 24 | 6 (25%) | 4 (16%) |

We analysed deceased (P0) and E19.5 mutant embryos and found large holes in their diaphragms (Fig1A-G). Younger mutant embryos (E14.5 and E16.5) were also found to have diaphragmatic holes (Fig1H-J and Fig1K-R), often accompanied by liver herniation into the thoracic cavity (Fig1I, J, L, and M). 80-90% of CDH is Bochdalek-type, characterised by hernias in the posterolateral region of the diaphragm. In > 85% of the cases, Bochdalek hernias are left-sided^14^. A description of the phenotypes of the *Prx1^Cre/+^;Wt1^loxp/loxp^* embryos at different stages is summarised in Table 2, with the left sided hernia being the most common defect (45%). 20% of the mutants had bilateral hernias, and 5% had holes on the right side only. Finally, 30% of the mutants either did not exhibit any obvious diaphragmatic phenotype or did not have fully formed holes but with thinning of the diaphragm was observed.

**Figure 1.**
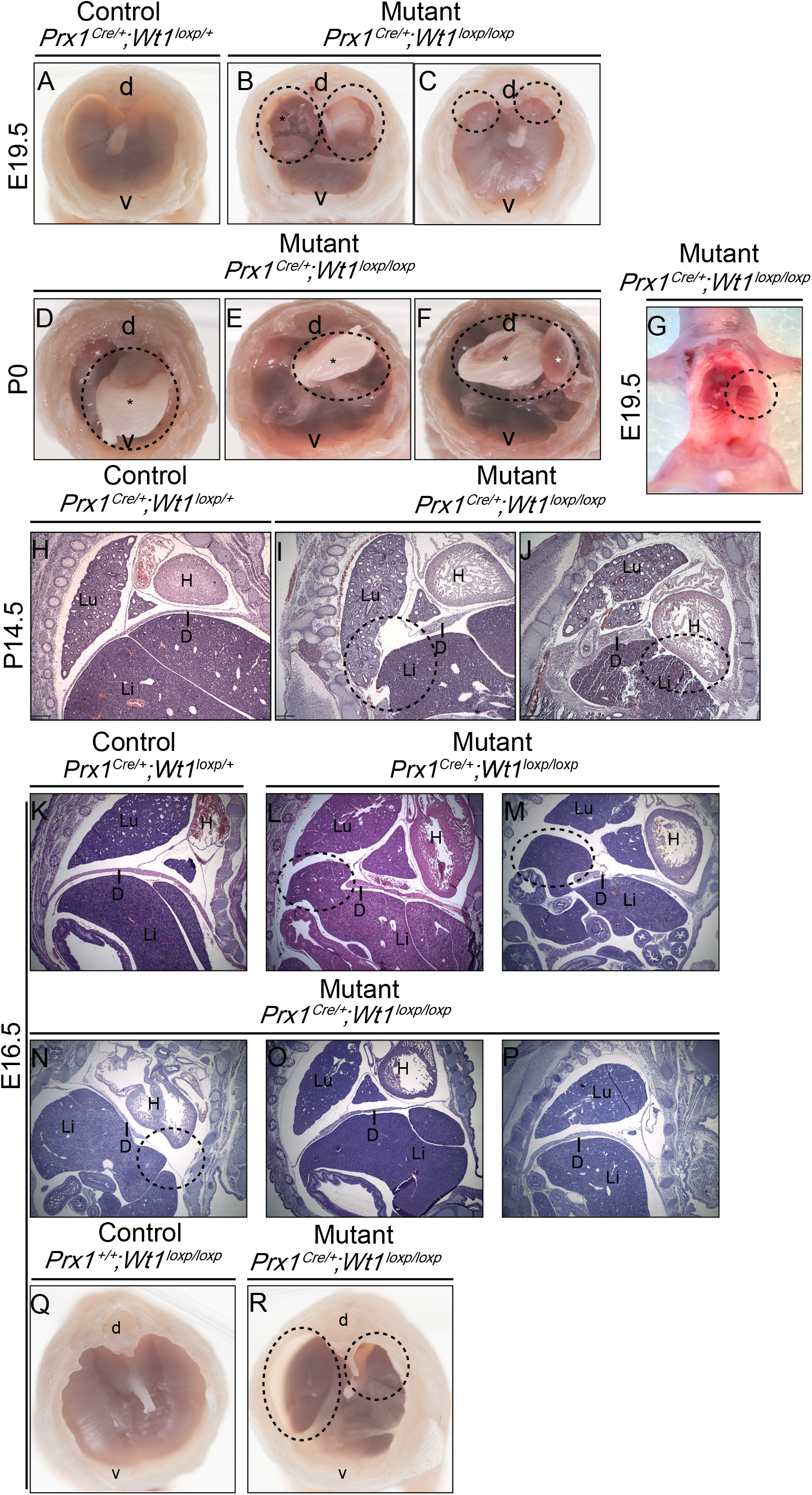
Deletion of *Wt1* using Prx1Cre leads to mutant embryos and pups with diaphragmatic hernia (*Prx1^Cre/+,^ Wt1^loxp/loxp^*) Male *Prx1^Cre/+;^Wt1^loxp/+^* mice were crossed with females *Wt1^loxp/loxp^*. Embryos from various stages or pups (which died shortly after being born) were taken for analysis. Mutants (*Prx1^Cre/+;^Wt1l^oxp/loxp^*) exhibited diaphragmatic hernia. (**A-G**) Pups were dissected, heads removed and thoracic content removed, and the diaphragm imaged from the top. (**B,C**) Images of hernias in the E19.5 mutant embryos. (**B**) Bilateral and dorsally located holes were found in the diaphragm (circled), with liver herniation into the thoracic cavity (asterisk). (**C**) Smaller bilateral and dorsally located holes were observed (circled) and litter mate control is shown in (**A**). (**D-F**) Representative images of hernias in the P0 mutant pups. Large and left-sided dorsally located holes (circled) with herniation of the stomach (black asterisk) and liver herniation (white asterisk). (**G**) Image of an E19.5 mutant embryos imaged from below with abdominal content removed, revealing a left-sided dorsally located hole in the diaphragm (circled). E14.5 (**H-J**) and E16.5 embryos (**K-P**) were sectioned and H&E stained. Holes in the diaphragm are circled. (**Q,R**) Freshly isolated E16.5 embryos were dissected, contents of thoracic cavity removed, and imaged from the top of the embryo. (**R**) Bilateral dorsal holes were present in the mutant embryo (circled). (Abbreviations: d: dorsal, v: ventral, Lu: lung, H: heart, Li: liver, D: diaphragm)

**Table 2.** Phenotype descriptions for the (Prx1^Cre/+^;Wt1^loxP/GFP^) embryos and pups at different developmental stages

| EMBRYO STAGE | TOTAL # MUTANTS | GENDER/DIAPHRAGM PHENOTYPES |
| --- | --- | --- |
| E14.5 | 3 | Hole left side, dorsal, with liver herniation.<br>Hole left side, ventral, with liver herniation.<br>Hole in right side dorsal, possible liver herniation. |
| E16.5 | 8 | Female. Bilateral holes, dorsal, with liver herniation.<br>Male. Hole left side, dorsal, with liver herniation.<br>Female. No obvious hole/hernia – possible thinning of diaphragm on left side. NB: Embryo sectioned in the coronal plane.<br>Male. No obvious hole/hernia. Possible thinning at edges.<br>Male. No obvious hole/hernia.<br>Female. Bilateral holes, dorsal, with liver herniation.<br>Female. No obvious hole/hernia.<br>Male. No obvious hole/hernia. |
| E18.5 | 1 | Bilateral thinning of the diaphragm, dorsal. No obvious hole/herniation. |
| E19.5 | 4 | Bilateral holes, dorsal, with liver herniation.<br>Small bilateral holes/thinning of the diaphragm, dorsal.<br>Hole left side, dorsal.<br>Hole left side, dorsal. |
| P0 | 4 | Male. Large hole left side, dorsal, with stomach and liver herniation.<br>Male. Large hole left side, dorsal, with stomach and liver herniation.<br>Female. Large hole left side, dorsal, with stomach and liver herniation.<br>Female. Large hole left side, dorsal, with stomach and liver herniation. |

The initial separation of the thoracic and peritoneal cavities is established by the PPFs, which are typically formed between E10.5-E12.5^5,4^. By E12.5 a continuous band of cells has developed, completely separating the two cavities. H&E stained sections of E11.5 and E12.5 mutant embryos (Fig2 A-D’, sectioned transversely) show that this continuous band of cells failed to form correctly (Fig2 B, B’, D,D’). Large gaps between the PPFs and the PMPH were observed in the mutants (Fig2 B, B’, D, D’), leading to the failure of separation of the two cavities.

**Figure 2.**
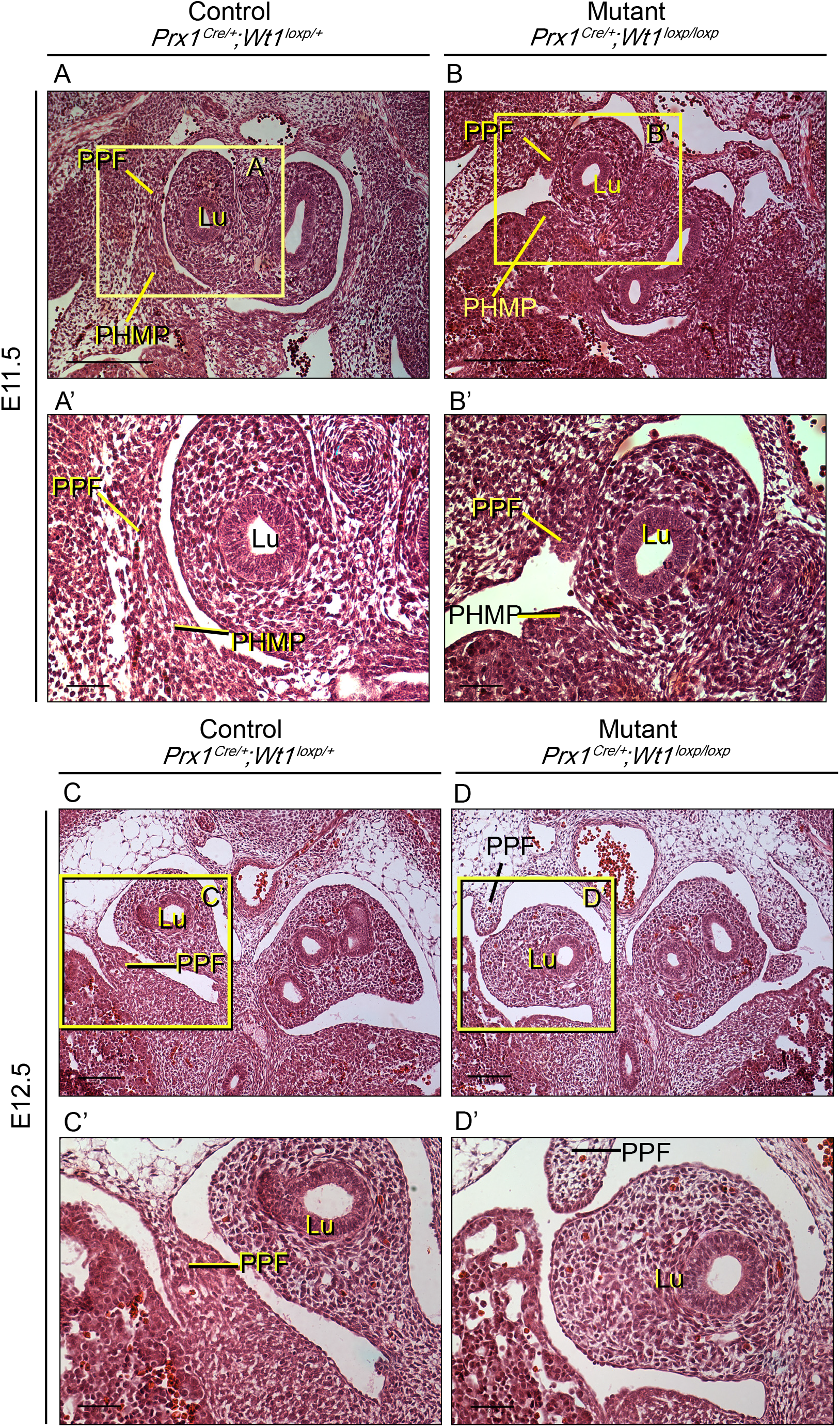
E11.5 and E12.5 *Prx1^Cre/+^;Wt1^GFP/loxP^ embryos* have disrupted PPF formation. (**A,B**) Representative Images of E11.5 H&E stained sections are shown, illustrating disrupted PPF development in the mutants (**B**). A gap is visible between PPF and PHMP. A litter mate control is shown in (**A**). **A’** and **B’** are magnified images of the areas boxed in **A** and **B**. (**C,D**) Representative images of E12.5 H&E stained sections are shown. Images of disrupted PPF development is shown in (**D**) and images of litter mate control showing a continuous band of PPF/PHMP is shown in (**C**). **C’** and **D’** are magnified images of the areas boxed in **C** and **D**. *(Abbreviations: Lu, Lungs; PPF, pleuroperitoneal fold; PHMP, posthepatic mesenchymal plate)*. (Scale bar = 100 μm in A, B, C, and D; 50 μm in A’, B’, C’, and D’).

### Prx1-Cre labelling of the Wt1+ mesenchymal cells in the PPFs

To understand the cause of this defect we first needed to know the location of Prx1-Cre expression (i.e. in which cells *Wt1* was being deleted). Immunofluorescence was performed on sections from *Prx1^Cre/+;^R26R^mTmG/+^* lineage tracing embryos using anti-GFP and anti-WT1 antibodies. In this model, cells in which Prx1-Cre is or has been expressed are GFP positive. As shown in Fig3, WT1 expression was detected in the PHMP and PPFs (indicated in red). However, whilst the cells of the PPFs expressed GFP and were thus Prx1-Cre lineage positive (indicated in green), no expression of GFP was detected in the cells of the PHMP (Fig 3 A, A’). This was particularly obvious in the E11.5 embryos. At E11.5, the PPFs and PMHP have just begun to fuse, with boundaries still clearly visible between the two structures. This boundary is distinctly marked by the clear domains of GFP expressing cells. To better demonstrate this, consecutive sections in this region were obtained and stained (Fig3 A and B, where B is anterior to A). In A’ and A’’, the PPFs (GFP+) and PHMP (GFP-) can be seen abutting one another, whilst in B’ and B’’, it is clear that the PPFs (GFP+) are migrating downwards and infiltrating the PHMP (GFP-). At E12.5, once the continuous band has been formed, the population of GFP+ cells (PPF derived) infiltrating the PHMP is still distinctive (Fig 3C). As for other visceral organs, the diaphragm is lined by a mesothelial layer. The mesothelial cells of the diaphragm express endogenous WT1 (as indicated by the yellow arrowhead in Fig 3C’); however, this mesothelial layer does not express GFP and therefore is not derived from the Prx1-Cre lineage and thus not from the PPFs. Using this model we have demonstrated that only the PPFs arise from Prx1-Cre expressing cells and not the PHMP. Therefore, despite WT1 being expressed in both the PPFs and PHMP, it will only be deleted in the cells of the PPFs in our model.

**Figure 3.**
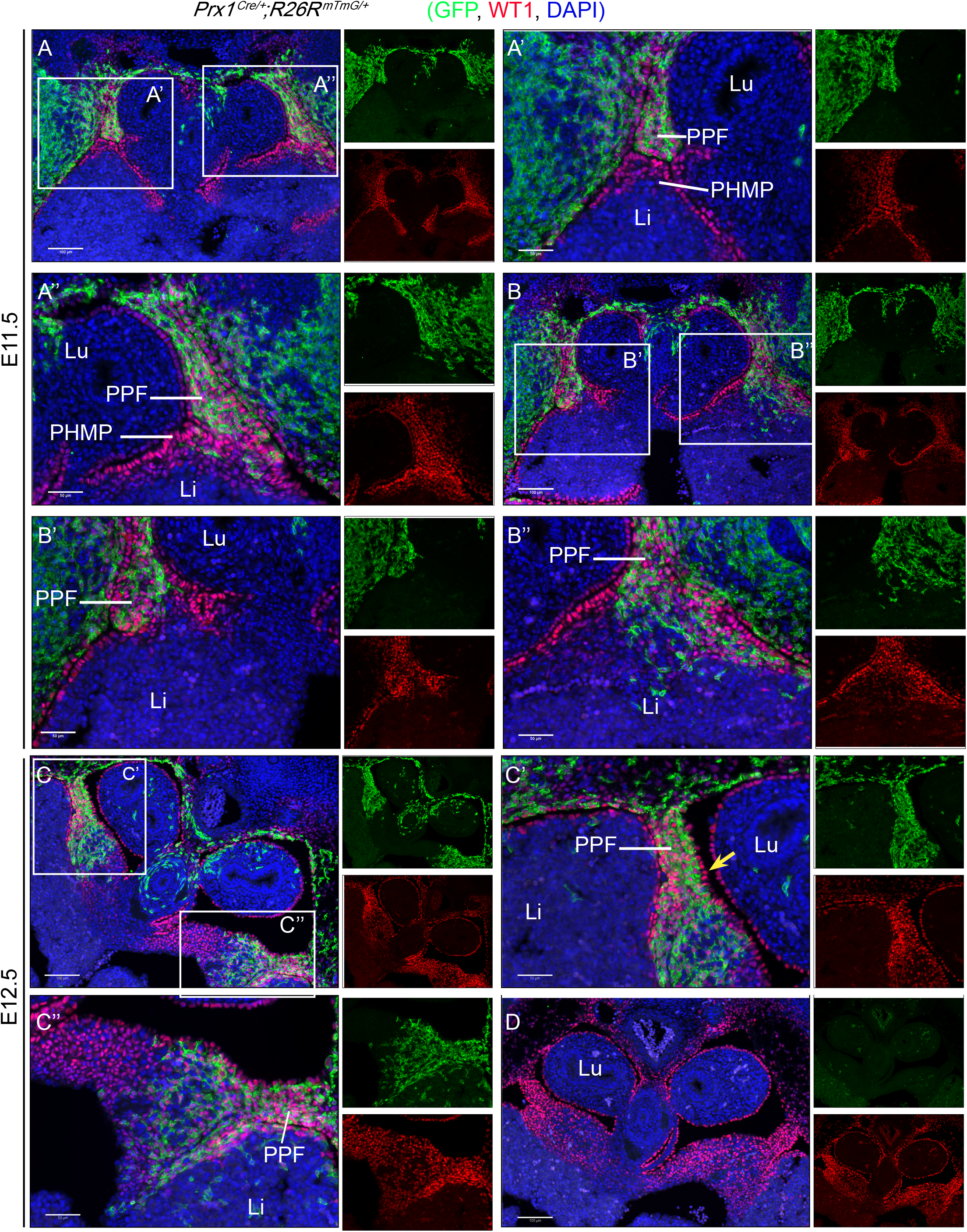
The PPFs but not the PHMPs, are Prx1-Cre lineage positive. Transversely sectioned E11.5 or E12.5 Prx1-Cre lineage tracing embryos (*Prx1^Cre/+^;R26R^mTmG/+^*) are stained with an anti-GFP antibody (indicated in green) and an anti-Wt1 antibody (red). Cell nuclei are stained with DAPI (blue). **A** and **B** are consecutive sections, where **B** is anterior to **A**. Wt1 expression (red) is detected in both PPFs and PHMPs, while GFP signal is only detected in PPFs. B” indicates the migration of PPFs into the PHMPs starting to form a continuum band. (**C**) This PPF/PHMP continuum is fully formed at E12.5. GFP expressing cells are located at the edge where the PPF and PHMP are merged/fused. The mesothelial cells of PPFs did not express GFP (yellow arrow in C’). **A’, A”, B’, B”, C’**, and **C’’** are magnified images of the areas in **A, B,** and **C**. A Prx1Cre negative litter mate control is shown in (**D**). *(Abbreviations: Lu, lungs; Li, liver; PPF, pleuroperitoneal fold; PHMP, posthepatic mesenchymal plate)*. (Scale bar = 100 μm in A, B, C, and D; 50 μm in A’, A’’, B’, B”, C’, and C’’). (n=3).

To better illustrate this, a second mouse model was generated. *Prx1^Cre/+^;Wt1^GFP/+^* males were crossed with *Wt1^loxP/loxP^* females. In *Prx1^Cre/+;^Wt1^loxP/GFP^* offspring, one copy of the *Wt1* allele is flanked by loxP sites whilst the other copy has a GFP knockin at exon 1 which disrupts function. Therefore, Wt1 expressing cells will be GFP positive. Moreover, upon Cre-mediated loxP recombination (driven by Prx1-Cre), the second copy of *Wt1* is conditionally deleted. The mutant cells remain GFP positive despite no functional WT1 being present. Staining with anti-GFP and anti-WT1 antibodies, reveals cells that have once expressed Wt1 but no longer do so. Such cells will be positive for GFP but negative for endogenous WT1. This is illustrated in Fig4 (E11.5, E12.5 and E13.5), where endogenous WT1 expression (red) was detected in PPFs and PHMP in the control embryos (Fig 4A, A’, C), but its expression was completely lacking in the PPFs and unaffected in the PHMP of the mutant embryos (Fig 4B, B’, D, F). GFP expression was observed in the disrupted PPFs thus confirming that WT1 would normally be expressed in these cells (Fig 4B, 4B’, D, E, and F). Intriguingly, the GFP signal in the PPFs of the mutant embryos was much stronger than that in the control littermates (Fig 4). It is plausible that this is a consequence of an accumulation of cells in the PPFs due to defective cell migration. Identifying PPF and PHMP structures is not trivial as sections have to be obtained at precisely the same level from control and mutant embryos. This model provides a reassuring way of identifying the regions in which *Wt1* has been deleted, hence convenient for subsequent studies of pathways that are potentially disrupted.

**Figure 4.**
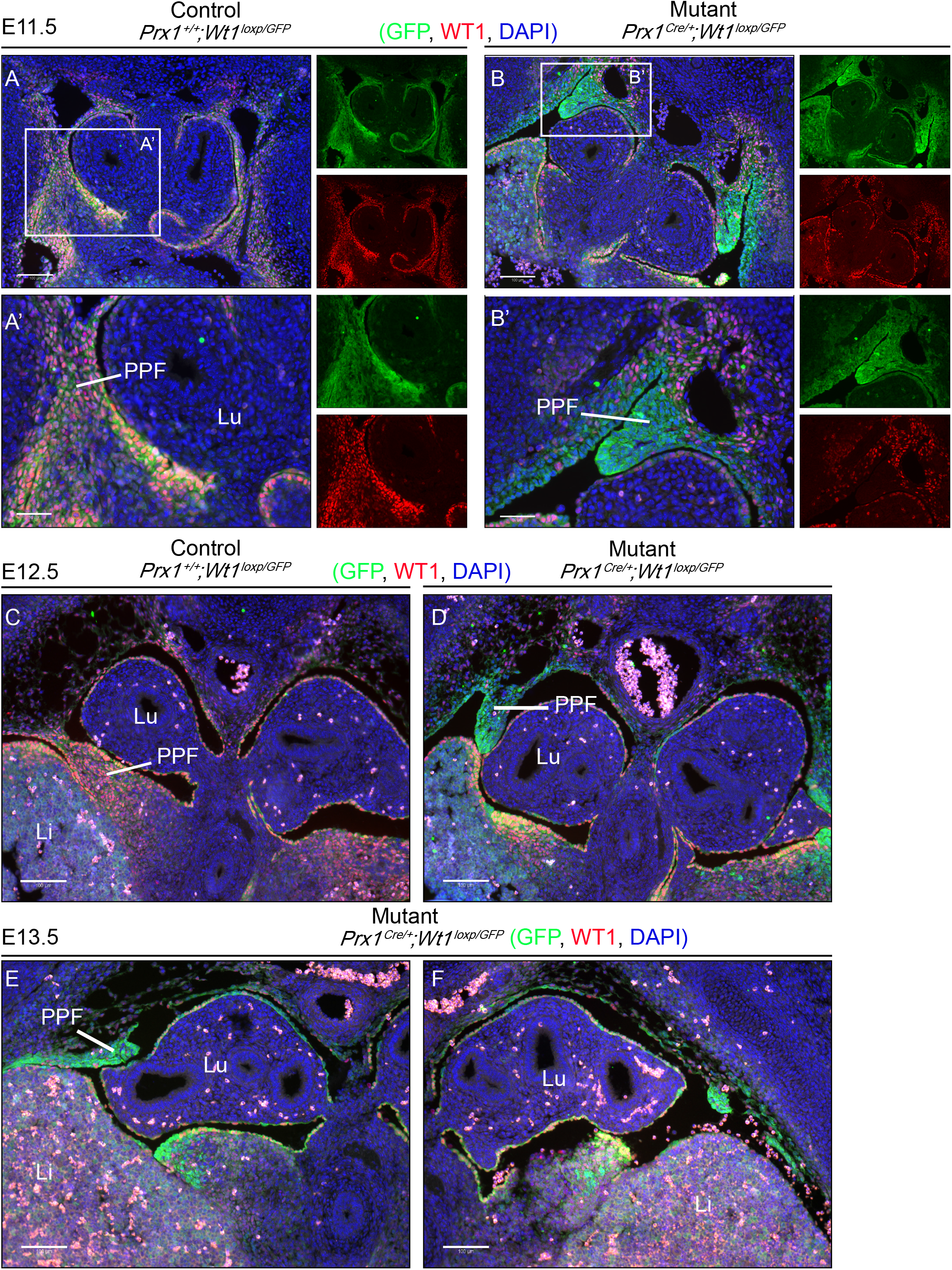
Wt1 deletion in the PPFs leads to disrupted PPF development in the *Prx1^Cre/+;^Wt1^GFP/loxP^* embryos, where GFP expression indicates cells that have had Wt1 deletion. Representative images of E11.5 (**A, A’, B, B’**), E12.5 (**C,D**), and E13.5 (**E,F**) sectioned embryos. Slides were stained with an anti-GFP antibody (green-upper inset), an anti-Wt1 antibody (red-lower inset), and DAPI (blue). Images from mutant embryos are shown in **B, D, E, and F**, where disrupted PPF expressed GFP but not Wt1, indicating deletion in this structure. A Prx1Cre negative littermate control from each stage is shown in **A, and C. A’**, **B’** are magnified images of areas indicated in **A and B**. *(Abbreviations: Lu, lungs; Li, liver; PPF, pleuroperitoneal fold; PHMP, posthepatic mesenchymal plate)*. (Scale bar = 100 μm in A, B, C, D, E, F; 50 μm in A’ and B’). (n=3).

### The epithelial status of PPFs deleted for Wt1 is unchanged

Previously we have shown that Wt1 plays a major role in regulating the epithelial and mesenchymal cell states in several mesodermal tissues^15^. To test the hypothesis that the formation of the continuous band of cells is disrupted due to defects in PPF cell migration, we analysed the expression levels of EMT markers. We stained sections from control (*Prx1^+/+^;Wt1^GFP/loxP^*) and mutant (*Prx1^Cre/+^;Wt1^GFP/loxP^*) embryos with an anti-E-cadherin (CDH1, an epithelial marker) antibody and an anti-Vimentin (VIM, a mesenchymal marker) antibody. Sections were co-stained with an anti-WT1 antibody or anti-GFP antibody to demonstrate deletion of Wt1 and regions of PPFs and PHMP. Despite strong expression of CDH1 in the lung bud (a positive control), no expression of CDH1 was detected in the PPFs of the mutant or control embryos (Supp Fig 1 CD’, CDH1 is indicated in green and GFP is indicated in red). Therefore, we did not observe any change in CDH1 expression in the PPFs between the control and mutant embryos. We also compared expression of vimentin between control and mutant embryos (vimentin is indicated in green and WT1 is indicated in red); however, no changes were observed (Supp Fig 1A-B’). Next we investigated whether cell proliferation within the mutant PPFs was reduced. However, staining with an anti-Ki67 antibody revealed no differences between control and mutant embryos. Therefore, the lack of a continuous band of cells is not caused by a disruption in PPF cell proliferation. (Supp Fig 2, Ki67 is indicated in red and GFP is indicated in green).

The retinoic acid (RA) signalling pathway plays a key role in diaphragm development. A commonly used mouse model of CDH utilises nitrofen to inhibit RA synthesis^16,17^. We analysed the expression of RALDH2, which catalyses the formation of RA from retinaldehyde. Importantly, WT1 has been shown to transcriptionally activate *Raldh2* in epicardial cells^18^. Surprisingly, we observed an increase in RALDH2 expression in the PPFs of the mutant embryos (Supp Fig 3B, B’).

To understand plausible molecular pathways that might be underlying the defects in our model, we generated a mouse model to allow the precise isolation of cells in the diaphragmatic region in which *Wt1* has been deleted (*Prx1^Cre/+^;R26R^tdRFP/+;^Wt1^loxP/GFP^*). In this model, cells that are derived from the Prx1-Cre lineage and have had *Wt1* deleted will express GFP and RFP. We used fluorescence-activated cell sorting (FACS) to sort for these cells from the diaphragmatic region of E11.5 embryos. Control cells were sorted from the same region isolated from a comparable model, differing only by the absence of loxP sites flanking one of the Wt1 alleles (*Prx1^Cre/+^;R26R^tdRFP/+;^Wt1^+/GFP^*) (therefore avoiding the deletion of *Wt1* in the Prx1-Cre lineage). We measured levels of expression of known EMT markers in these cells, including snai1 and snai2. Supp Fig 4 shows that the levels of these EMT markers are not altered, which is consistent with the E-cadherin and Vimentin results described in Supp Fig 1.

### PPFs give rise to non-muscle mesenchyme. Muscle connective tissue fibroblasts and myoblasts are absent in mutant PPFs

Defects in the PPFs have been suggested to be the cause of CDH in several mouse models^5^. The PPFs have been shown to give rise to the central tendon as well as the muscle connective tissue fibroblasts^5^. In addition, work performed by Paris *et al*. shows that WT1 expressing cells in the PPFs are non-muscle mesenchymal cells and majority of them do not express TCF4^12^. Together, these suggest that the TCF4-expressing cells (muscle connective tissue fibroblasts) and the WT1-expressing cells of the PPFs are likely to be two distinct cell types. In addition to the central tendon and the muscle connective tissue fibroblasts, our data suggest that the PPFs also give rise to the non-muscle mesenchyme (marked by WT1). We examined the expression of TCF4 in our model in which *Wt1* is deleted in the non-muscle mesenchymal cells by Prx1-Cre. Unexpectedly, TCF4 expression (indicated in red) was almost completely absent in the mutant PPFs at both E12.5 and E13.5 (Fig 5A-E). This result was not expected as Wt1 is not expressed in the muscle connective tissue fibroblasts in PPFs, suggesting possible interactions between these two populations of cells.

**Figure 5.**
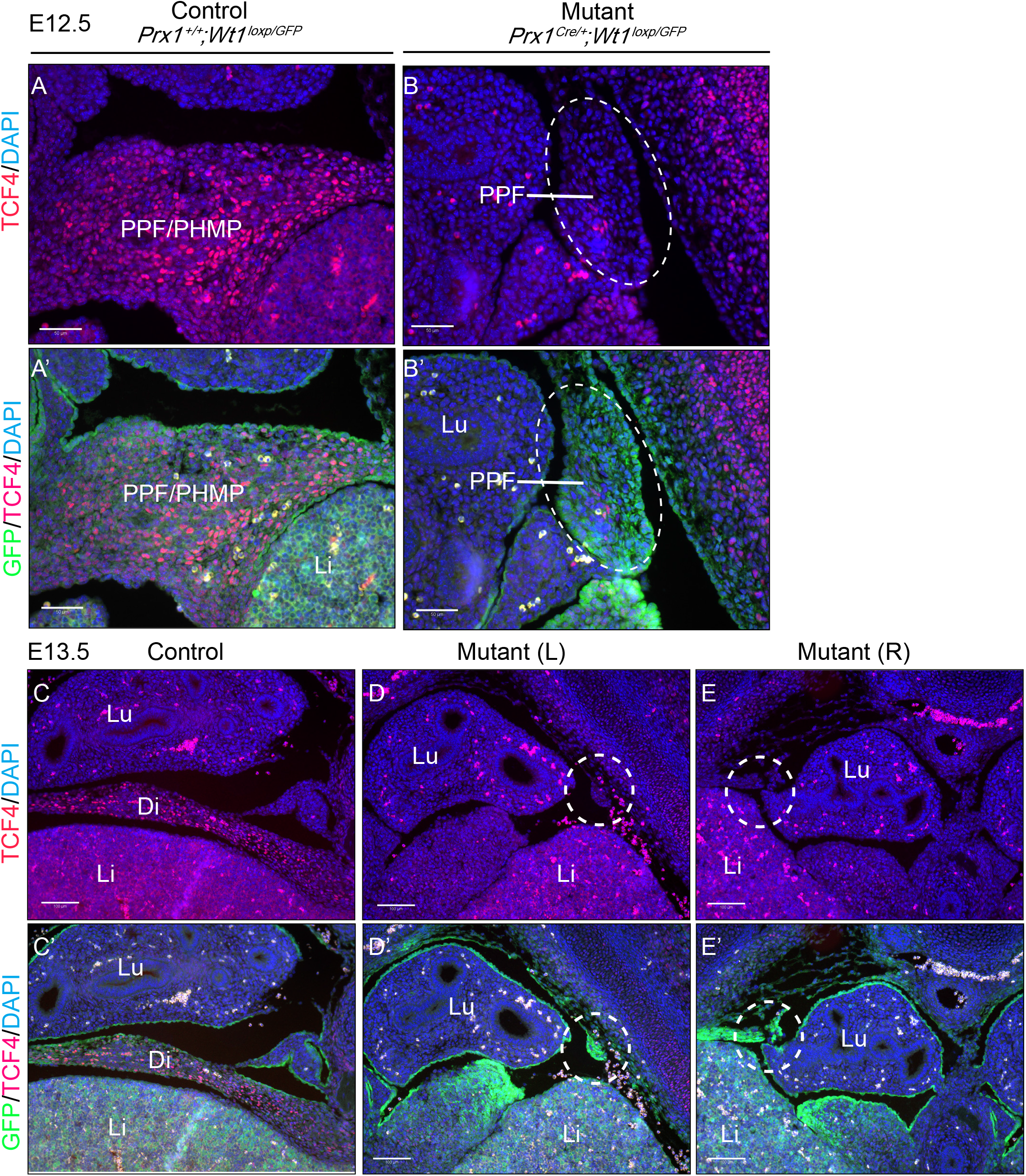
Wt1 deletion in the PPFs leads to absence of TCF4-expressing cells in the PPFs of *Prx1^Cre/+;^Wt1^GFP/loxP^* embryos. Representative images of E12.5 and E13.5 embryos were sectioned and stained with an anti-TCF4 antibody (red), anti-GFP antibody (green), and DAPI (blue). At E12.5 (**A, A’, B, B’**), TCF4 expressing cells are located in the PPF/PHMP continuum in the control embryo (**A, A’**) while these cells are absent in the mutant embryos (**B, B’**). At E13.5, TCF4 expressing cells (red) are located in the diaphragm of the control embryos (**C, C’**) and absent in the disrupted PPFs/PHMPs in the mutant embryos (**D, D’, E, E’**). Left side of disrupted PPF/PHMP is shown in (**D, D’**) and right side is shown in (**E, E’**). **A’, B’, C’, D’**, and **E’** are magnified images in **A-E**. *(Abbreviations: Lu, lungs; Li, liver; Di, diaphragm; PPF, pleuroperitoneal fold; PHMP, posthepatic mesenchymal plate)*. (Scale bar = 50 μm in A, A’, B, B’; 50 μm C, C’, D, D’, E, E’).

As well as labelling muscle connective tissue fibroblasts, TCF4 is a transcription factor that binds to B-catenin. The Wnt/B-catenin signalling pathway is known to act downstream of WT1^19^. The reduction in TCF4 expression in the mutant PPFs in our model could therefore be due to the disruption of the Wnt/B-catenin pathway as a result of reduced WT1 expression. Alternatively, the disappearance/absence of muscle connective tissue fibroblasts, which are marked by TCF4, could also explain the reduction. To test the second possibility, sections were stained with an anti-GATA4 antibody which also marks muscle connective tissue fibroblasts^5^. At E12.5, GATA4 expression is found at high levels in cells in of the PHMP as well as in the continuous band of cells separating the cavities (Fig 6A, A’). The expression is particularly strong in the mesothelium of the PPFs but much weaker (if not absent) in the mesenchymal/fibroblast cells within the PPFs (Fig 6A, A’). As with the TCF4 staining, no GATA4 expressing cells were found in the PPFs in which Wt1 was deleted (Fig 6B, B’, C, C’), E12.5 embryos from the *Prx1^Cre/+^;Wt1^loxP/GFP^* model where Gata4 is indicated in red and Wt1 and/or GFP is indicated in green). We also checked GATA4 expression at E13.5. Similarly, no GATA4 expression was detected in the rudiment of the PPFs in the mutant embryos (Fig 6E, E’, F, F’) compared with clear expression in the controls (Fig 6D, D’), suggesting an absence of muscle connective tissue fibroblasts in the mutant embryos.

**Figure 6.**
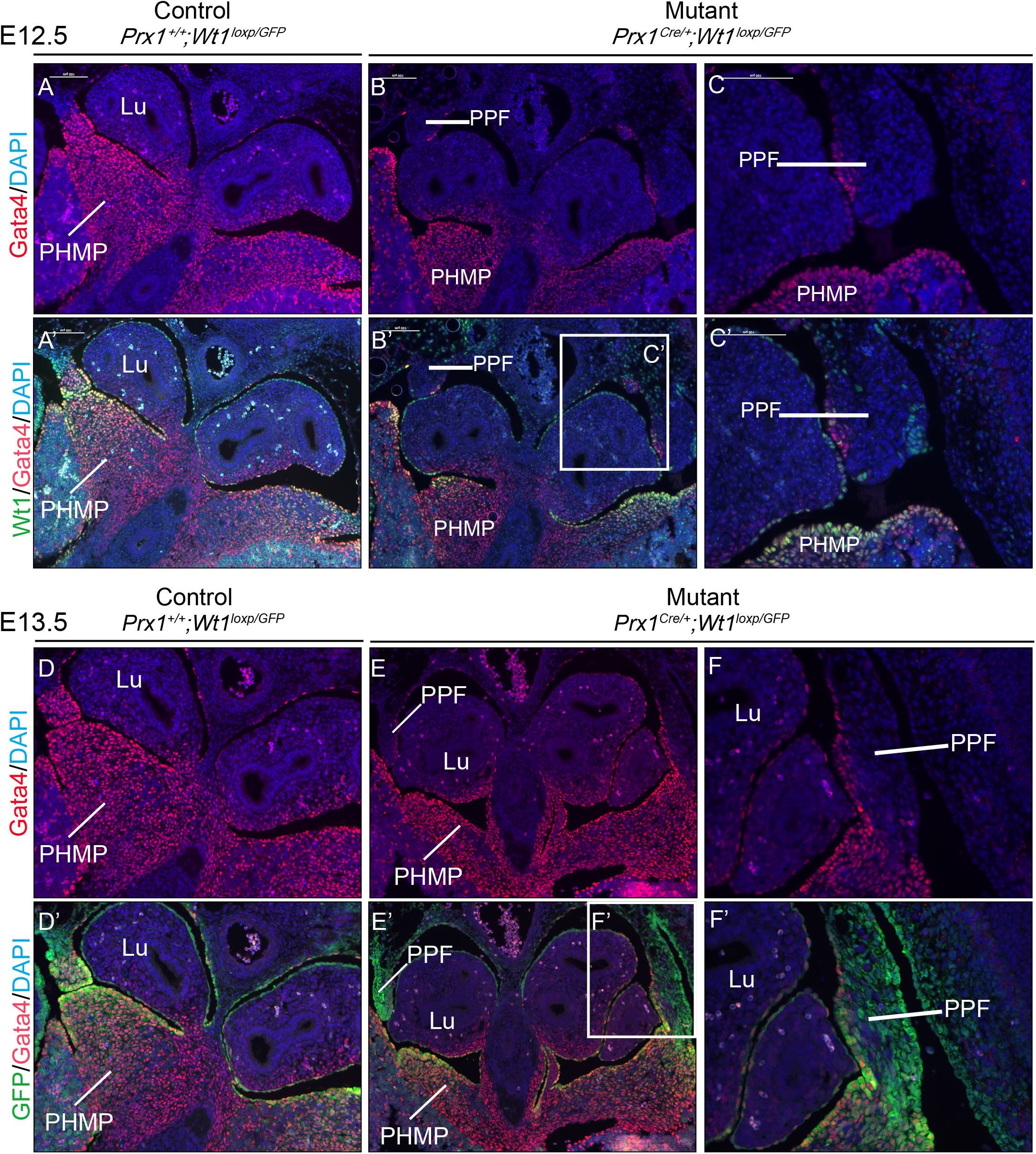
Wt1 deletion in the PPFs leads to absence of Gata4-expressing cells in the PPFs of *Prx1^Cre/+;^Wt1^GFP/loxP^* embryos. In **(A, A’, B, B’, C, C’**), representative images of E12.5 embryos were sectioned and stained with an anti-GATA4 antibody (red), anti-Wt1 antibody (green), and DAPI (blue). Images from mutant embryos are shown in (**B, B’, C, C’**), where Wt1 expression in the PPF is deleted and the Gata4-expressing cells are absent in the PPFs. Images from a Prx1-Cre negative littermate control are shown in (**A, A’**). Magnified areas indicated in **B** are shown in **C** and **C**’. (n=2 for controls and mutants). In **(D, D’, E, E’, F, F’**), representative images of E13.5 embryos were sectioned and stained with an anti-GATA4 antibody (red), anti-GFP antibody (green), and DAPI (blue). Representative images from Prx1Cre negative littermate controls are shown in (**D, D’**). Images of mutant embryos showing disrupted PPF development are shown in (**E,E’**). (**F,F’**) contain magnified images of indicated areas in (E, E’). *(Abbreviations: Lu, lungs; PPF, pleuroperitoneal fold; PHMP, posthepatic mesenchymal plate)*. (Scale bar = 50 μm in A, A’, B, B’; 50 μm C, C’, D, D’, E, E’). (Scale bar = 100 μm in A, A’, B, B’, D, D’, E, E’; 50 μm C, C’, F, F’).

The GATA4 and TCF4 expressing muscle connective tissue fibroblasts have been shown to play a crucial role in guiding the migration of myoblasts^5^. Myogenic progenitors begin to migrate from the somites to the diaphragm at around E12.5^3^. To test if the myoblasts were affected in our model, we stained the sections with an anti-MyoD antibody. In the control E12.5 diaphragm, MyoD positive cells are clearly seen (indicated in red, Fig 7A) whilst there is a complete absence of MyoD positive cells in the mutant PPF rudiments (Fig 7B, B’, C, C’). The myoblasts are present in the somites suggesting their formation is not affected (Fig 7B and C, myoblasts in somites are circled). In the mutant diaphragm at E16.5, there is muscle formation but the orientation of the muscle fibres appears to be disrupted, giving a “bunched up” appearance. Additionally, a thickening of the diaphragm is observed at the edges of the hernia, consistent with previous reports of disrupted migration in other models of CDH^20^. Sections are stained with an anti-MF20 antibody, which detects heavy chain of myosin II, indicated in green, and WT1 is indicated in red (Fig 7D, D’, E, E’). These data suggest that deleting *Wt1* in the non-muscle mesenchyme results in an absence of muscle connective tissue fibroblasts due to unknown mechanisms and a failure of myoblast migration to the PPFs, ultimately leading to disrupted diaphragm formation.

**Figure 7.**
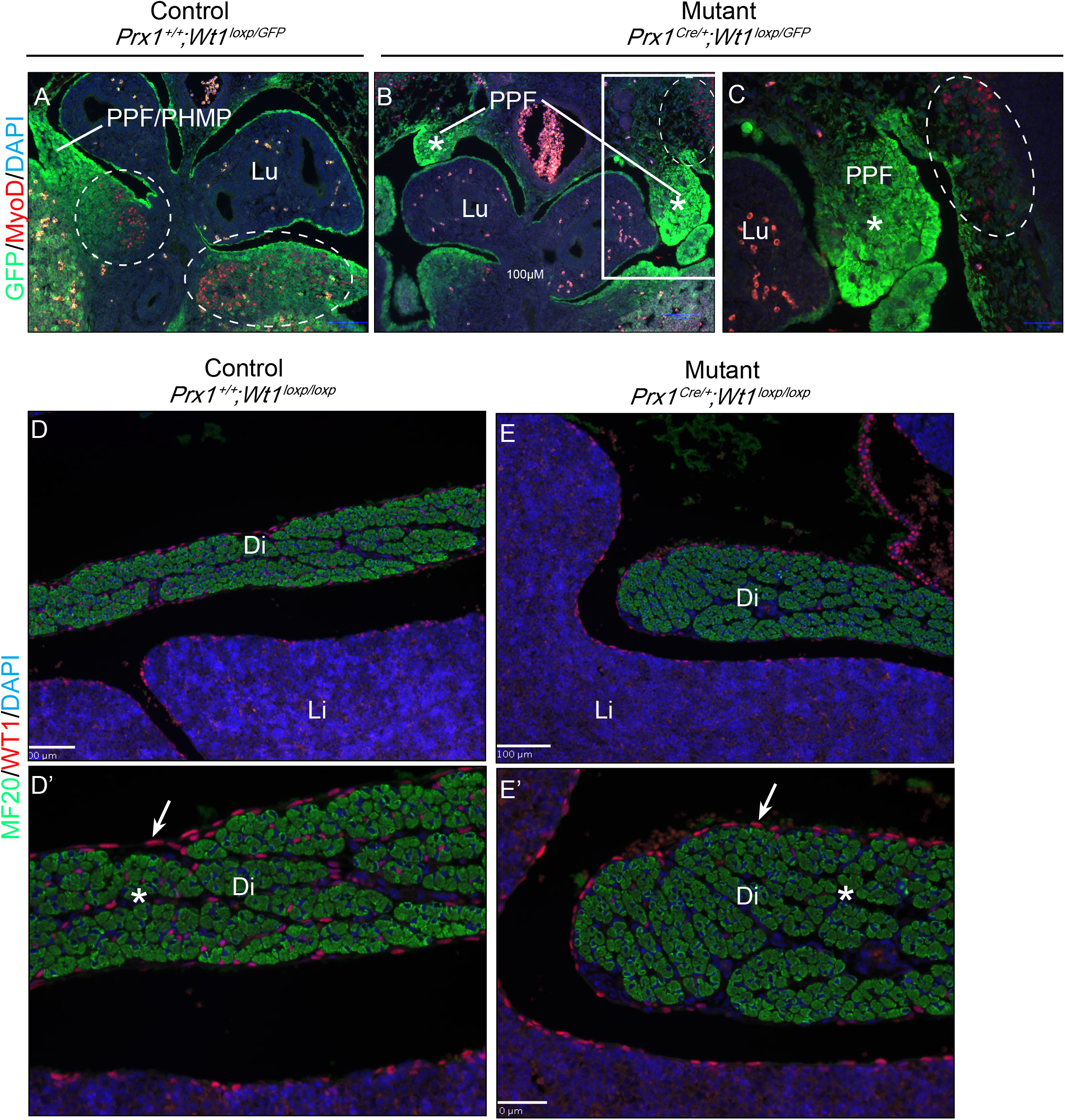
Wt1 deletion in the PPFs leads to absence of MyoD-expressing myoblasts in the PPFs of *Prxl^Cre/+;^Wt1^GFP/loxP^* embryos. Representative images of E12.5 embryos were sectioned and stained with an anti-MYOD antibody (red), anti-GFP antibody (green), and DAPI (blue). Images from control embryos, showing myoblasts (red, circled) located in the PPF/PHMP continuum (**A, A’**). In the mutant embryos, myoblasts are absent in the disrupted PPF/PHMP (**B, B’, C, C’,** white asterisks). Myoblasts are present in the somite (**B’**, **C’**, circled). Magnified areas in (**B,B’**) are shown in (**C, C’**). (D, D’, E, E’) Sagittal plan sectioned E16.5 embryos are stained with an anti-MF20 (green) and an anti-Wt1 antibody (red). The orientation of muscle fibres were mis-arranged (white arrow). WT1 expression is lost in the mesenchymal cells of the diaphragm (white asterisk) but not in the mesothelial cells (red arrow). *(Abbreviations: Lu, lungs; PPF, pleuroperitoneal fold; PHMP, posthepatic mesenchymal plate, Li: liver, Di: diaphragm)*. (Scale bar = 100 *μ*m in A, A’, B, B’; 50 μm C, C’).

## Discussion

CDH is a complex condition and many aspects of this disease are poorly defined, particularly the underlying causal mechanisms. The formation of diaphragm is a delicate process involving multiple cell types arising from several regions (somites, lateral plate mesoderm, neurons, and septum transversum). Different cell types interact with one another (for example muscle connective tissue fibroblasts guide the migration of the myogenic progenitors^5^) and crosstalk between signalling pathways is common. The complexity of diaphragm development no doubt contributes to the high frequency of CDH. Here we use transgenic mouse models to dissect the role of one crucial gene, *Wt1*, in the mesenchymal cells of the PPFs.

Firstly, we demonstrate the heterogeneous nature of the mesenchymal cell populations within the PPFs. As described previously, Prx1-Cre (expressed in the lateral plate mesoderm during development) gives rise to muscle connective tissue fibroblasts, marked by TCF4 and GATA4^5^. In agreement with a previous study^12^, we show that most of the TCF4+ (and GATA4+) cells in the PPFs do not express WT1. This suggests that in addition to the muscle connective tissue fibroblasts^5^, Prx1-Cre expressing cells give rise to an additional cell type in the developing diaphragm, with mesenchymal properties (marked by WT1). We next show that these WT1-expressing PPF cells (from the Prx1-Cre lineage) have the ability to migrate, and do so towards the PHMP, infiltrating it to form a continuous band of cells, ultimately sealing off the thoracic and peritoneal cavities. It is interesting to note that in the CDH model in which *Gata4* is deleted in the muscle connective tissue fibroblasts using Prx1-Cre (*Prx1Cre^Tg/+^;Gata4^∇/flx^;Rosa26^LacZ/+^*), B-gal+ sacs are present covering herniated regions, and not as holes in this tissue^5^. Since our data demonstrate that cells in the Prx1-Cre expressing lineage can give rise to mesenchymal cells which have the ability to migrate, it is likely that these mesenchymal cells might contribute to the thin membranous sac formed in the model described by Merrell *et al*^5^. However, in our model, in which *Wt1* is deleted in mesenchymal cells of the PPFs using Prx1-Cre, we observe a failure of closure of the body cavities at early stages and holes/hernia at later stages. Together, these suggest that the Wt1-expressing mesenchymal cells of the PPFs can be one driving force behind cell migration from the PPFs towards PHMP.

Moreover, our data suggest that cells of the PHMP may also migrate towards the PPFs. In the diaphragms of E12.5 embryos (at which point the continuous PPF/PHMP membrane is almost complete), GATA4 expression is high in the cells of PHMP and most of these GATA4+ cells co-express WT1. Our staining suggests that the GATA4+ cells in the PHMP migrate towards the PPFs, as most of the cells in the PPFs do not express GATA4. This forms a distinct boundary at the point at which the PPFs (GATA4-) and PHMP (GATA4+) meet. A similar staining pattern is described in a recent study where G2-Gata4-Cre is used to delete *Wt1*^4^.

Connective tissue fibroblasts (indicated in yellow in cartoon illustration, Fig 8) are absent from the PPFs in our model in which *Wt1* is deleted in the non-muscle mesenchymal cells (indicated in green in cartoon illustration, Fig 8) using Prx1-Cre. Interactions between two cell types are common. For example, in skeletal muscle, ablation of muscle satellite cells (Pax7+) severely reduces the expansion of muscle connective tissue fibroblasts (TCF4+) during regeneration^21^. The molecular mechanisms that might govern the interactions between these cell types during diaphragm development remain incompletely understood (cartoon illustration, Fig 8).

**Figure 8.**
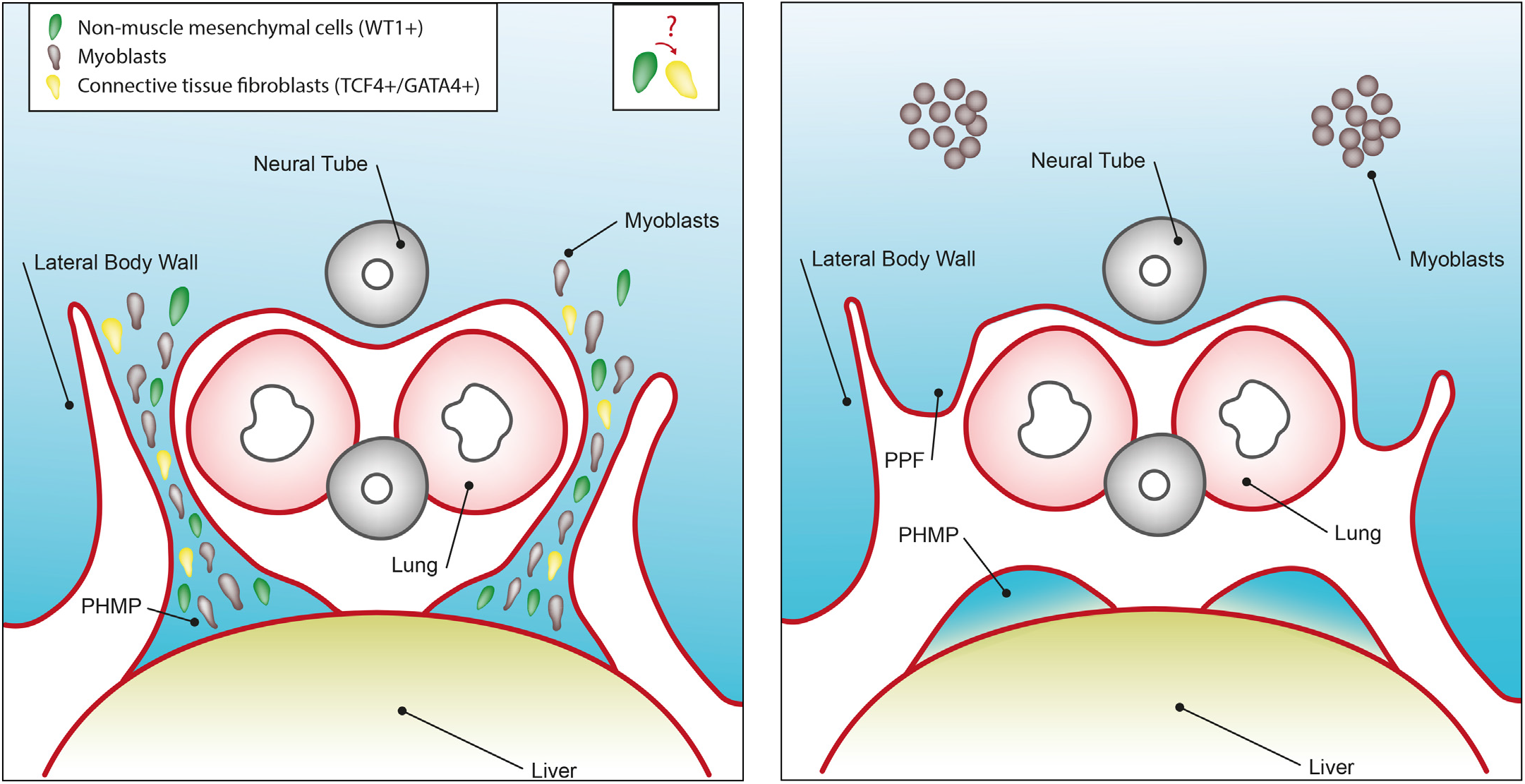
A cartoon illustration of diaphragm formation. Data from our model suggest the heterogeneity of mesenchymal cells in the PPFs and the importance of the Wt1-expressing non-muscle mesenchymal for diaphragm development. (**A**) Normal and (**B**) mutant condition where *Wt1* is conditional deleted by a Prx1-Cre. We demonstrate the non-muscle mesenchymal cells (indicated in green) in the PPFs migrate into PHMPs and form a continuous band. Deletion of *Wt1* using Prx1-Cre leads to failure of these mesenchymal cells moving towards PHMPs and form a gap. In the PPFs deleted for *Wt1*, connective tissue fibroblasts (indicated in yellow) are absent for unknown reasons. We speculate a possible cross talk between non-muscle mesenchymal cells and the connective tissue fibroblasts. Absence of connective tissue fibroblasts may lead to the disruption of myoblasts (indicated in brown) migrating to the PPFs.

In addition to connective tissue fibroblasts (yellow, Fig 8) and non-muscle mesenchymal cells (green, Fig 8), there is an additional player in diaphragm development, myoblasts (brown, Fig 8). Absence of connective fibroblasts has previously been shown to cause defects in guiding the myogenic progenitors towards PPFs^5^. Staining with a muscle progenitor marker, myoD, confirmed this is indeed the case in our model. It should be noted that muscle fibres are present in the mutant diaphragm at later stages (e.g. E16.5), with possible disrupted orientation. In addition, shorter and thicker segments of diaphragm are present adjacent to the hernia in our mutant embryos at later stages. It is possible that muscle progenitors still retain some ability to migrate into the PPFs (or ‘ride on’ other moving cells as the embryo grows and expands in size), but less effectively, in the absence of the guiding connective tissue fibroblasts.

These data lead us to investigate the underlying molecular mechanisms of the paused PPFs. Defects in EMT would be a plausible cause; however, we do not see changes in EMT markers at the protein (e.g. E-Cadherin and Vimentin by immunostaining) or mRNA (e.g. *snai1* and *snai2* by QPCR) level. A recent study elegantly showed that conditionally deleting *Wt1* using a G2-Gata4-Cre results in the mice developing diaphragmatic hernias. Defects in EMT (epithelial to mesenchymal transition as indicated in their study by an increase in E-cadherin, an epithelial cell marker), were suggested to be the underlying cause^4^. The Cre activity of G2-Gata4-Cre is detected in PHMP but in only some of the PPFs at E10.5 and E11.5^4^. Whilst in our model we show that cells of the Prx1-Cre lineage are retained in PPFs and start to migrate from PPFs towards PHMP. The difference between these results might be due to the different mouse models used, here we delete *Wt1* in PPFs using Prx1-Cre, and the model described by Carmona *et al*. results in the deletion of W*t1* predominantly in the PHMP. It is plausible that the cells that express Wt1 in the PPFs differ from those that reside in the PHMP, as the two structures are derived from different regions during development.

Interestingly, we observed an increase in RALDH2 expression in the mutant PPFs deleted for *Wt1* in our model. This unexpected increase in RALDH2 in diaphragms deleted for Wt1 has also been described in work performed by Carmona *et al*^4^. In their model, they reasoned that the increase in RALDH2 was due to the intermediate mesoderm of the renal ridges persisting in the PPFs. In situ hybridization of *Pax2* (a marker of the intermediate mesoderm) provided evidence for their argument. However, no PAX2 expression was observed in the PPFs in our model (using an anti-PAX2 antibody, Supp Fig 3C). Clear PAX2 staining was observed in the neural tube, suggesting the antibody worked well (Supp Fig 3D). Again, the difference between these results may be due to *Wt1* deletion using different Cre lines, hence affecting different regions or structures.

In summary, we illustrate the heterogeneous nature of the cell population that forms the PPFs, using several novel mouse models. We show that the conditional deletion of *Wt1* in the non-muscle mesenchyme using Prx1-Cre results in diaphragmatic hernias, and illustrate that the cells of the PPFs have the ability to migrate towards PHMP. Connective tissue fibroblasts are absent in the PPFs in which the non-muscle mesenchyme is deleted for *Wt1*, suggesting possible interactions between these two cell types. This absence of connective tissue fibroblasts could be a plausible explanation for the evident disruption to the migration; however, more work is required to delineate the exact underlying molecular mechanism.

## Materials and Methods

### Mouse husbandry

Animals used in this study were housed at the animal facilities at the University of Edinburgh with procedures performed under Personal and Project Home Office Licenses. Targeted deletion of *Wt1* (*Prx1^Cre/+^;Wt1^flx/flx^*) was achieved by crossing homozygous mice^15^ carrying loxP-flanked *Wt1* allele into a mouse strain of cre recombinase expression driven by the Prx1 transgene^24^. The Wt1-GFP mouse line (*Wt1^GFP/+^*) used in this study was made by Hosen *et al*^25^. GFP is a knockin at the first exon of *Wt1* and is expressed under the endogenous transcriptional regulatory elements of *Wt1*. The Prx1-Cre lineage tracing model (*Prx1^Cre/+^; R26R^mTmG/+^*) was made by crossing male Prx1-Cre mice with female *R26R^mTmG/mTmG^* double fluorescence reporter mice^26^. Cre expression, following Prx1 derived enhancer element, mediates a switch of Tomato to eGFP expression. The model to obtain mutant diaphragmatic cells, that are expressing Wt1 and have been conditionally deleted for *Wt1* by the Prx1-Cre (*Prx1^Cre/+;^ W1^GFP/flx^; R26R^tdRFP/+^*), are generated by crossing *Prx1^Cre/+^; Wt1^GFP/+^; R26R^tdRFP/tdRFP^* male with female *Wt1^flx/flx^*. To obtain counterpart control cells in the above model, male *Prx1^Cre/+^;Wt1^GFP/+^; R26R^tdRFP/tdRFP^* is crossed with wild type females.

### Tissue preparation

Samples were fixed in 4% paraformaldehyde (dissolved in PBS) at 4°C overnight, unless otherwise stated. Next day, samples were washed three times in PBS (10 min each) and stored in 70% ethanol before preparing for paraffin embedding. Tissue-Tek VIP^®^ Jr. Vacuum Infiltration Processor was used for paraffin wax embedding. Paraffin sections were cut 5-6 um thick using a microtome, mounted on SuperFrost^®^ Plus Microscope slides and dried at 50°C overnight.

### Immunofluorescence

Immunofluorescence staining was performed using a similar protocol as described previously^27^. The primary antibodies used in this study are Wt1 (Abcam ab89901, 1:1000), Ki67 (Abcam ab15580, 1:1000), GFP (Abcam ab5450, 1:1000), Pax2 (Biolegend 901001, 1:100), MF20 (DSHB Ab_2147781, 1:20), Vimentin (Santa Cruz sc-7557, 1:100), E-cadherin (BD 610181, 1:100), Raldh2 (Santa Cruz sc-22591, 1:200), Gata4 (Santa Cruz sc-25310, 1:100), Tcf4 (Cell Signalling C48H11, 1:100), and MyoD (Santa Cruz sc-32758, 1:100). Alexa-Fluor 488 or 594 conjugated antibodies were used as secondary antibodies. Sections were stained with DAPI and mounted using VectorShield. A Zeiss Axioplan II microscope was used to view immunofluorescence and H&E stained sections. Image capture was performed using the open source microscopy software: uManager.

### Haematoxylin and Eosin staining (H&E)

Slides were dewaxed in xylene, rehydrated in a series of ethanol washes, followed by washing in tap water, and stained with Mayer’s Haematoxylin. After washing with tap water, sections were differentiated in 1% HCl in 70% ethanol for a few seconds and washed with tap water. Slides were stained with saturated lithium chloride solution for a few seconds before washing in tap water. Slides were then stained with eosin, rinsed in tap water, and rehydrated and mounted using DPX mounting medium.

### Flow cytometry and fluorescence activated cell sorting (FACS)

Tissues of thoracic region/upper abdominal region (with lung, heart, and liver removed), were digested into single cell suspension using collagenase (1mg/ml collagenase and 4mg/ml BSA, dissolved in PBS) for 30-60 minutes at 37°C with shaking. Collagenase activity was stopped by washing the cells in PBS containing 5% FCS. Cells were pelleted by centrifugation at 300g for 5 min. Cells were filtered using 40 um cell strainer and subjected to FACS (BD FACSAriaTM II System). RNA from the sorted cells was extracted using TRIzol® Reagent.

### QPCR

cDNA was synthesised from RNA using a QuantiTect^®^ Whole Transcriptome kit (Qiagen) following the manufacturer’s protocol. QPCR was performed using the Universal Probe Library system (Roche) using LightCycler 480 II machine. The cycling conditions are preincubation (10 min, 95°C), amplification (95°C for 10 seconds, 60°C for 30 seconds, 72°C for 1 second), and cooling (40°C for 30 seconds). The amplification is repeated 50 times. Primer and probe information for each gene used is listed in Supp Table1.

---

**Supplementary Figure 1.**
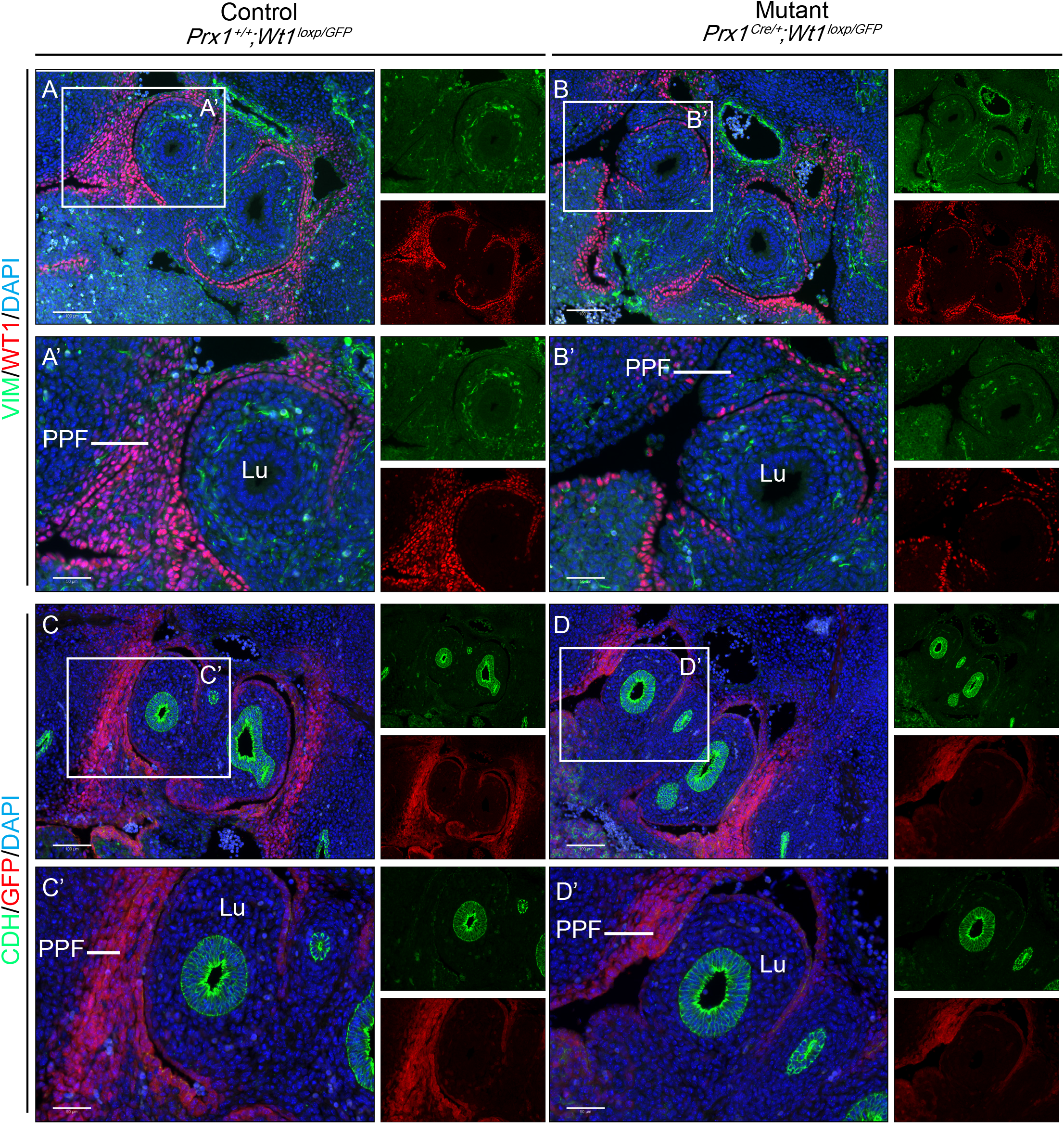

**Supplementary Figure 2.**
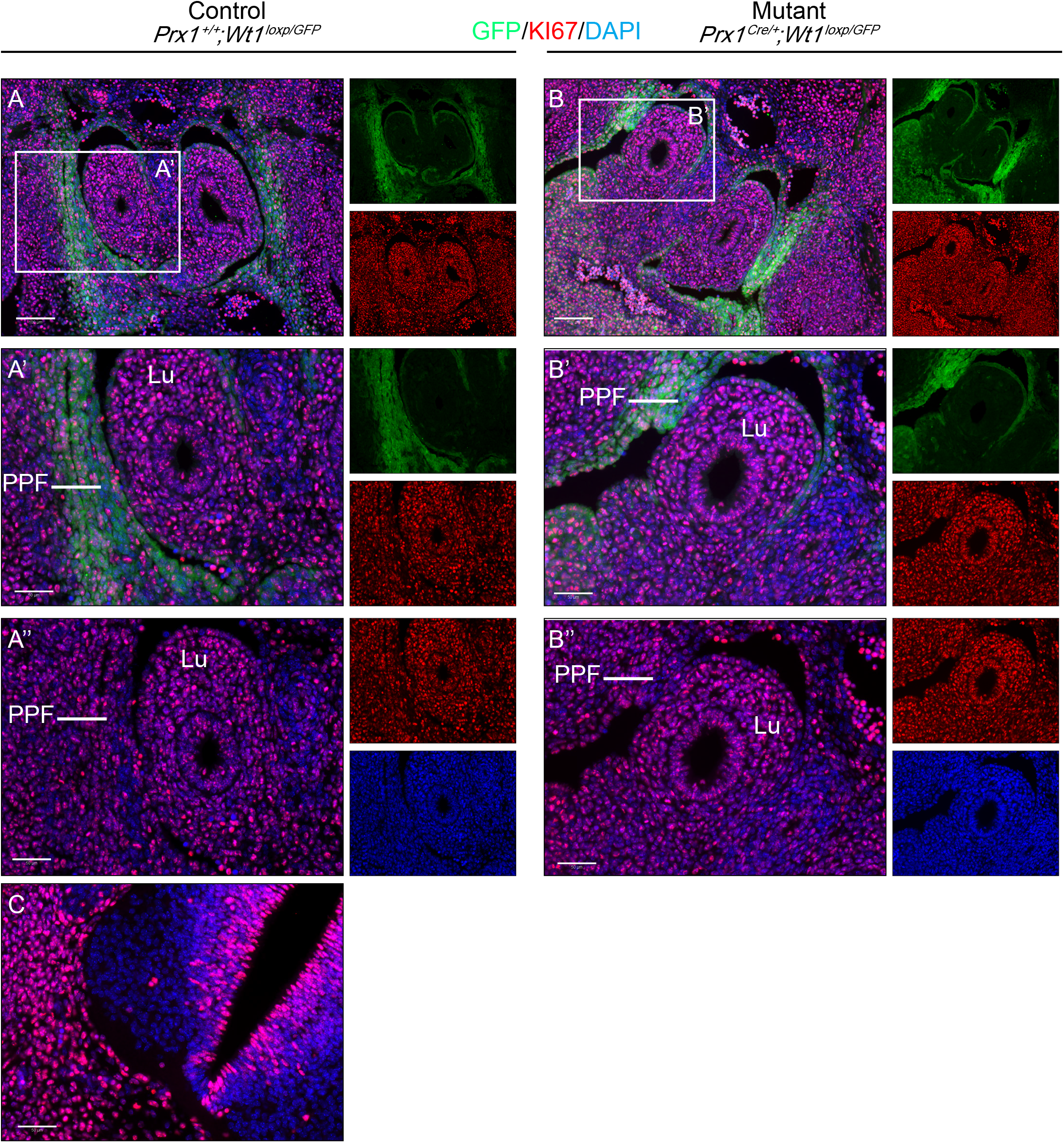

**Supplementary Figure 3.**
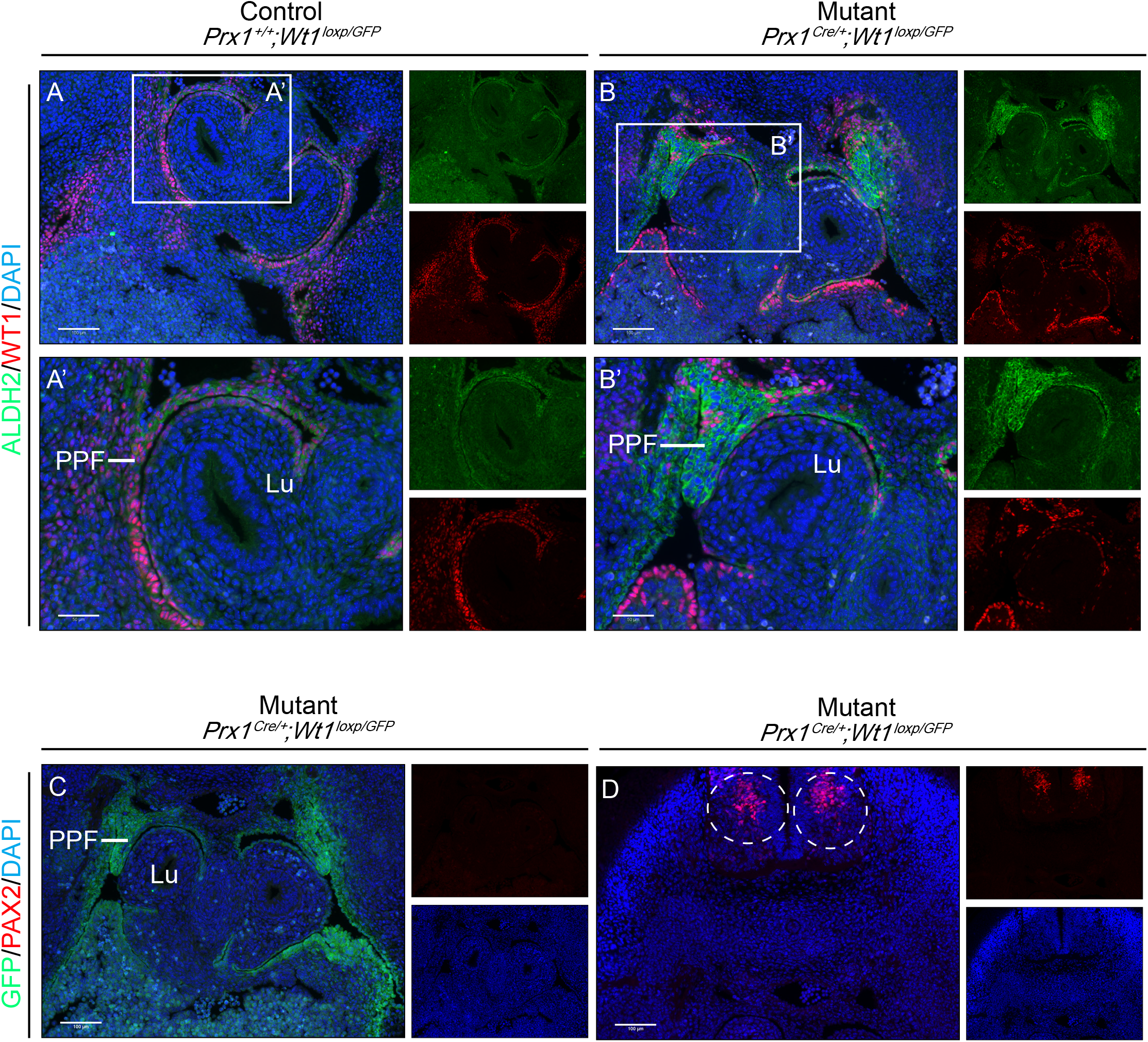

**Supplementary Figure 4.**
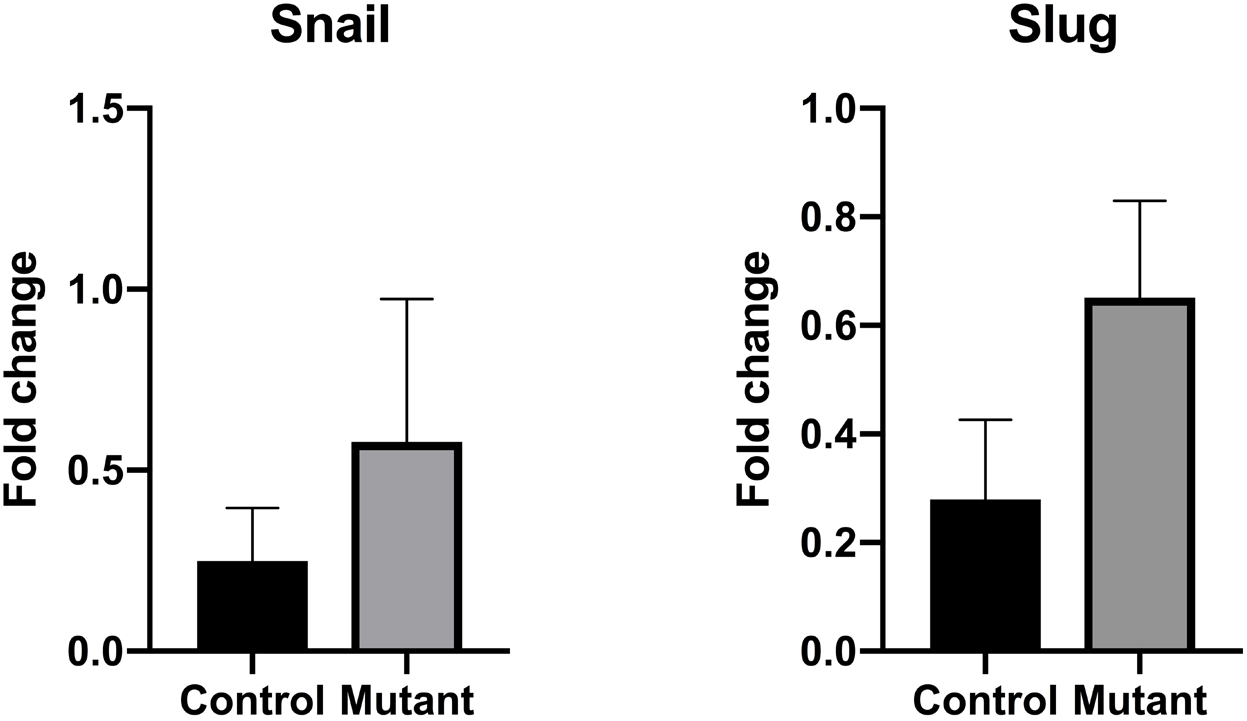

